# Microtubule-based mitotic spindles contain a micron-sized mixed-nucleotide zone

**DOI:** 10.1101/2021.07.23.453504

**Authors:** Cédric Castrogiovanni, Alessio Inchingolo, Jonathan U. Harrison, Damian Dudka, Onur Sen, Nigel Burroughs, Andrew D. McAinsh, Patrick Meraldi

**Author notes:** Equal contribution (Cédric Castrogiovanni, Alessio Inchingolo).

## Abstract

Current models infer that the microtubule-based mitotic spindle is built from GDP-tubulin with small GTP caps at microtubule plus-ends, including those that attach to kinetochores (K-fibres). Here we reveal that K-fibres additionally contain a dynamic mixed-nucleotide zone that reaches several microns in length. This zone becomes visible in cells expressing fluorescently labelled EBs, a known marker for GTP-tubulin, and endogenously-labelled HURP - a protein which we show to preferentially bind the GDP microtubule lattice *in vitro*. In living cells HURP accumulates on the ends of depolymerising K-fibres, whilst avoiding recruitment to nascent polymerising K-fibres. This gives rise to a growing “HURP-gap” which we can recapitulate in a minimal computational simulation. We therefore postulate that the K-fibre lattice contains a dynamic, micron-sized mixed-nucleotide zone.

**One Sentence Summary:** We reveal that the microtubules of the mitotic spindle contain a third, uncharacterized domain, a mixed nucleotide zone that resides between the GTP-cap and the GDP-tubulin lattice.

## Report

The mitotic spindle ensures faithful chromosome segregation during cell division. One of its key components are kinetochore fibres (K-fibres), bundles of ~15 microtubules that bind chromosomes via kinetochores, multiprotein complexes assembled on centromeres (*1*, *2*). By maintaining attachment to polymerising and depolymerising K-fibres, the kinetochores can generate the forces necessary for chromosome movement (*3*). The dynamicity of the microtubules depends on the equilibrium between GTP-cap growth, which occurs via addition of new GTP-tubulin dimers, and GTP-cap loss, which triggers depolymerisation of the GDP-tubulin lattice (*4*, *5*). Current models of the mitotic spindle imply that K-fibres are largely comprised of a GDP-lattice with the exception of a GTP-cap at the tip of polymerising kinetochore-bound microtubules (see model in **Fig. 1A**)(*6*). Based on structural data the GDP-tubulin lattice and GTP-tubulin enriched cap are thought to have different conformational states that are coupled to the nucleotide hydrolysis cycle (*7*). The End-Binding (EB) proteins, EB1 and EB3, read-out the location and size of the GTP-cap by preferentially binding the GTP-tubulin conformation (*8*). Moreover, *in vitro* experiments indicate that EB proteins also bind a transition form of αβ-tubulin, in which the GTP is already hydrolysed but the γ-phosphate group has not yet been released (*9*). *In vivo* EBs are limited to a narrow region of up to 100 nm, close to the kinetochore consistent with the presence of a small GTP-cap (*10*, *11*). Another microtubule-associated protein that binds to K-fibres in the vicinity of kinetochores is the hepatoma up-regulated protein (HURP) (*12*). Microtubule-binding by HURP is regulated by the RanGTP-gradient that forms around chromosomes (*13*), and displaces Importin-β from a microtubule-binding site on HURP. This leads to an accumulation of HURP on a region of the K-fibres proximal to kinetochore - termed “HURP stripes” (*12*, *14*, *15*).

**Fig. 1.**
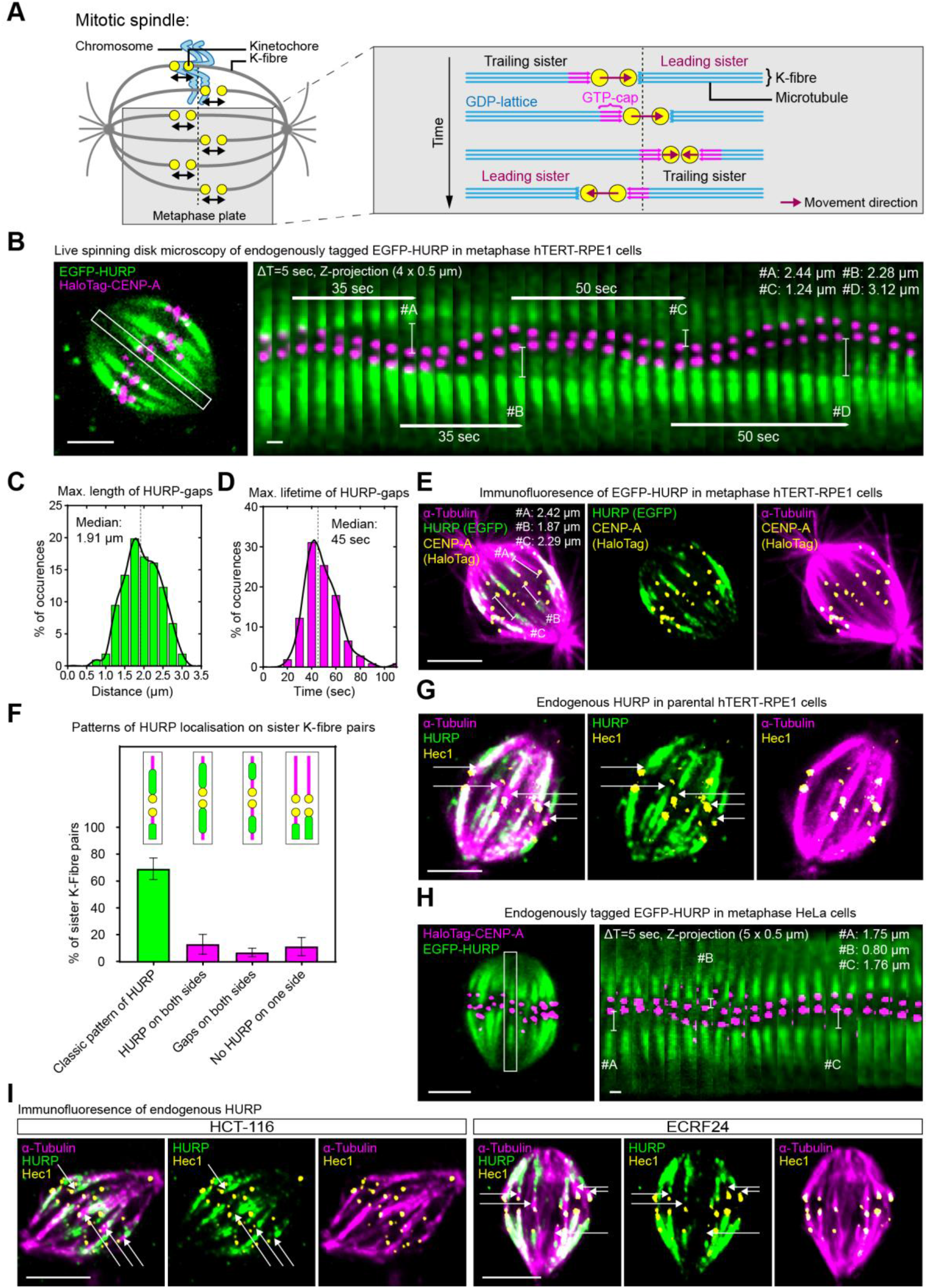
HURP is excluded from the K-fibre growth zone. (**A**) Current model for sister kinetochore oscillations in metaphase. (**B**) Live cell imaging of metaphase hTERT-RPE1 EGFP-HURP/HaloTag-CENP-A cell. Inset shows sister K-fibre pair used for kymograph (right panel), bars display the HURP-gaps maximum length. (**C** and **D**) Distribution of live HURP-gap maximum lengths (C) and duration (D; N = 4 independent experiments, n = 106 K-fibers in 29 cells) Black line = curve fit. (**E**) Immunofluorescence image of metaphase hTERT-RPE1 EGFP-HURP/HaloTag-CENP-A cell. Z-projection of 1 μm thickness. Bars show gap distances. (**F**) Quantification of mean HURP localization patterns along sister K-fibre pairs in fixed cells (error bars = s.d., N = 4, n = 419 sister K-fibres in 19 cells). (**G**) Immunofluorescence image of endogenous HURP in metaphase hTERT-RPE1 cell. Z-projection of 0.5 μm thickness. Arrows indicate HURP-gaps. (**H**) Live cell image of endogenously-tagged EGFP-HURP in metaphase HeLa cell overexpressing HaloTag-CENP-A. Inset shows sister K-fibre pair used for kymograph (right panel), bars display gap maximum distances. (**I**) Immunofluorescence images of endogenous HURP in metaphase HCT-116 and ECRF24 cells. Z-projection of 1.5 and 1.0 μm thickness, respectively. Arrows indicate HURP-gaps. Scale bars = 5 μm and 1 μm (kymograph). Gamma = 0.7 for EGFP-HURP-channel in E, G, and I.

When analysing the spatiotemporal dynamics of endogenously-tagged EGFP-HURP in human hTERT-RPE1 (non-transformed immortalized human retina pigment epithelial) cells expressing the kinetochore marker Halo-CENP-A, we observed a striking localization in metaphase that did not fit the expected Ran-GTP directed pattern: as sister-kinetochore pairs oscillated along the spindle axis, a wide gap formed on the growing K-fibres between the HURP stripes and the attached kinetochore; there was no such gap on the depolymerizing K-fibres, with EGFP-HURP conterminous with the kinetochore. (**Fig. 1B - MOVIE 1 & 2**). The median maximal HURP gap size on polymerizing K-fibres measured 1.91 μm with a lifetime of ~45 s (n = 106; **Fig. 1C & D**). Immunofluorescence staining with anti-GFP and α-tubulin antibodies confirmed that HURP-gaps were present on 49.6 % of metaphase K-fibres (median width of 0.94 μm; **Fig. 1E and Fig. S1A & S1B**), consistent with the notion that sister-kinetochores pairs are bound by one growing and one shrinking K-fibre. While most kinetochore pairs (69.2 ± 8%) displayed a HURP-gap on one side and a strong EGFP-HURP signal on the other side, 13 ± 7.4% were conterminous with HURP signal on both sides, and 6.8 ± 3.3% had HURP-gaps on both sides (**Fig. 1F and Fig. S1C**). These data are consistent with the fact that pairs can be transiently connected to shrinking or growing K-fibres on both sides (*16*). The HURP-gap was not a by-product of the GFP-tag or cell line since it was also visible after staining with antibodies directed against endogenous HURP (**Fig. 1G**) and visible in live images of HeLa (hypertriploid cervical carcinoma) cells expressing endogenously-tagged EGFP-HURP (**Fig. 1H & Movie 3**), as well as fixed HCT-116 (pseudo-diploid colorectal carcinoma) and ECRF24 (immortalized human umbilical vein endothelial) cells stained with HURP antibodies (**Fig. 1I**). These data suggest that in the metaphase spindle, HURP is absent from polymerizing K-fibre regions connected to kinetochores, despite high RanGTP concentrations.

To precisely define the relationship between HURP-gaps and kinetochore motion we used lattice-light sheet microscopy and kinetochore tracking to acquire and analyse 3D volumes of hTERT-RPE1 EGFP-HURP/Halo-CENP-A cells every 4.5 s (**Movie 4**) (*17*). As metaphase sister-kinetochores oscillated back-and-forth with K-fibres switching from growth to shrinkage, we found that HURP was exclusively present on depolymerizing K-fibres (associated to the leading kinetochore sister), and never on the growing K-fibre (associated to the trailing kinetochore sister) (**Fig. 2A & B**). As soon as a directional switch occurred, HURP started accumulating on the depolymerizing K-fibre that was previously devoid of HURP, and a gap started to form on the opposite, growing K-fibre (**Fig. 2B & C**). Higher HURP intensities on depolymerizing K-fibres correlated with a higher probability for a directional switch (**Fig. 2D**), suggesting that HURP may act as a microtubule-rescue factor. We conclude that HURP-gaps are associated with polymerizing K-fibres.

**Fig. 2.**
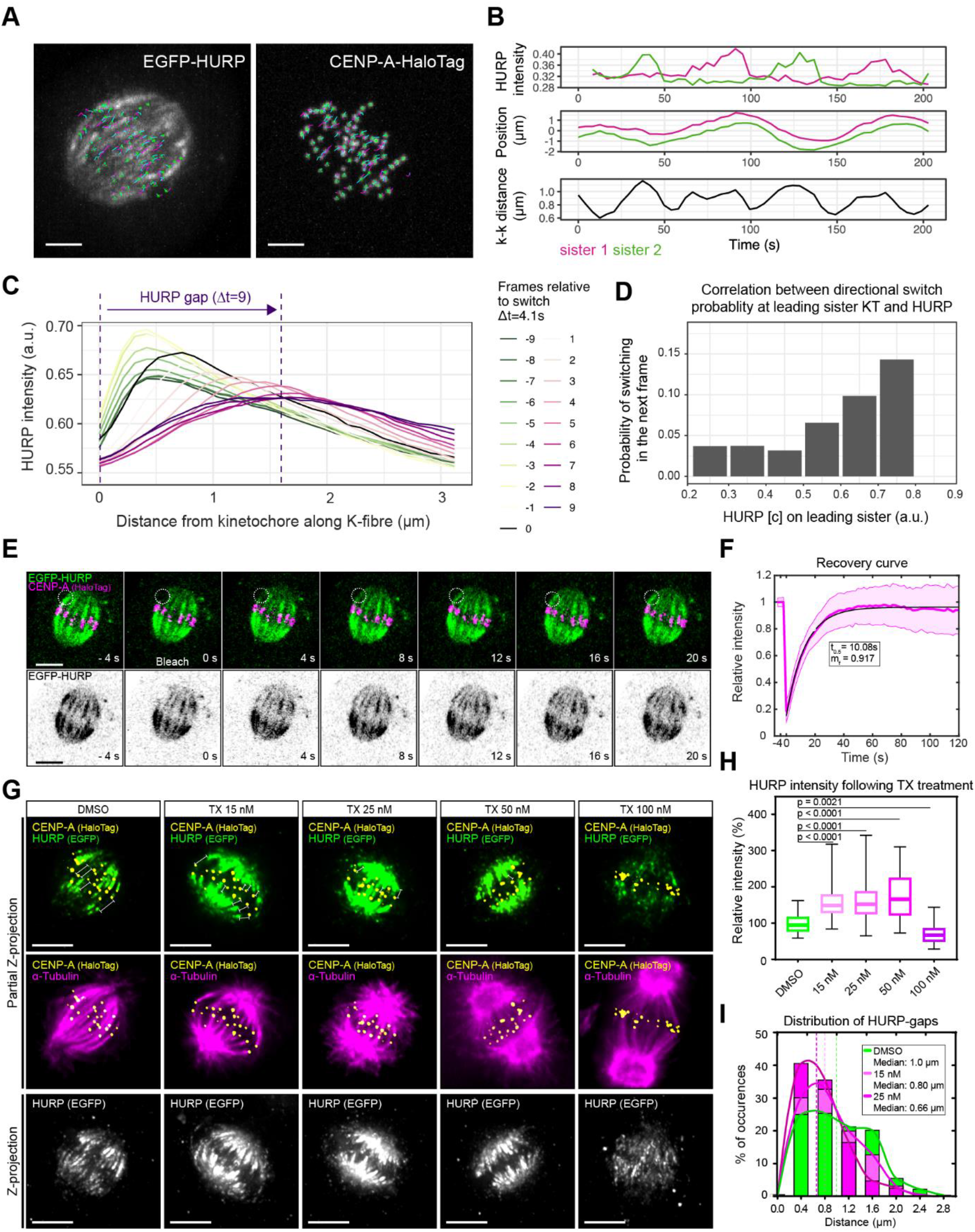
HURP-gaps are precisely linked to K-fibre conformation. (**A**) Lattice light sheet images of metaphase hTERT-RPE1-EGFP-HURP/HaloTag-CENP-A cell projected in Z. (**B**) Exemplary HURP intensities and sister kinetochore positions in single pair over time. (**C**) Average spatial distribution of HURP along K-fibre over time relative to switch from leading to trailing (N = 3, n = 69 pairs). (**D**) Probability of directional switch in the next frame versus HURP on the K.fibre in the current frame (N = 3, n = 69 pairs). (**E**) FRAP time-lapse of hTERT-RPE1 EGFP-HURP/HaloTag-CENP-A cell. Inset indicates bleached region on single K-fibre. (**F**) EGFP-HURP FRAP recovery curve. Error bars = s.d; Black line = fit of mono-exponential recovery (N = 4, n = 47 K-fibres). (**G**) Immunofluorescence images of metaphase hTERT-RPE1-EGFP-HURP/HaloTag-CENP-A cells treated with indicated taxol concentrations for 45 min. Gamma = 0.5 for GFP-channel; partial Z-projections of 1.0, 0.8, 0.9, 1.1 and 1.3 μm thickness, respectively. Bars show HURP-gap distances. Z-projection = projection of whole spindle. (**H**) Boxplots of relative HURP intensities in taxol-treated cells as shown in G (N = 4; n = 57-67 cells; p = Kruskal–Wallis test and Dunn’s multiple comparisons). (**I**) HURP-gap size distributions in fixed metaphase hTERT-RPE1-eGFP-HURP/HaloTag-CENP-A cells treated with DMSO, or 15 and 25 nM Taxol. (N = 4; n = 272-345 gaps). Lines = curve fit. Scale bars = 5 μm.

One possible reason for the formation of the HURP-gap is a slow on-rate for microtubule binding. However, fluorescence recovery after photobleaching (FRAP) experiments on single K-fibres in metaphase, revealed a fast on-rate (~10 s – mobile fraction 91.7 %) for EGFP-HURP (**Fig. 2E & F**). Because this value is 4-fold lower than the median HURP-gap lifetime (~45 s, **Fig. 1D**) we excluded slow binding as the cause of the HURP-gaps. A second possibility was that HURP can differentiate between “new lattice”, which is built from GTP-tubulin dimers, and the “old” GDP-tubulin lattice that forms as GTP is hydrolysed. To evaluate this possibility, we treated cells with increasing doses of the microtubule-stabilizing agent taxol. While at low concentrations taxol is thought to mainly suppress microtubule-ends dynamics, at higher concentrations it has been postulated to stabilize microtubules and induce a GTP-tubulin-like conformation in the microtubule lattice (*18*–*24*). Consistently, kinetochore-tracking and immunofluorescence experiments indicated that increasing taxol doses progressively led to rigid and straight K-fibres with decreasing chromosome movements (**Fig. 2G & S2A**). As long as chromosomes could still move, HURP intensities increased with increasing taxol concentrations (up to 50 nM), while the size of the HURP-gap shrunk (**Fig. 2G-I**). HURP levels, however, strongly decreased at the highest taxol concentration (100 nM), when chromosome movements were frozen (−31% vs DMSO; **Fig. 2G & H and Fig. S2A**). The general HURP decrease was not due to the loss of K-fibres since staining against the spindle checkpoint protein Mad2, a marker of unattached kinetochores (*25*), revealed few, if any, unattached kinetochores (**Fig. S2B**). This behaviour suggested that HURP association to microtubules increases as long as taxol only reduces plus-end dynamics, but decreases when taxol imposes a GTP-tubulin-like conformation on the microtubule lattice.

To investigate whether the *in vivo* behaviour of HURP can be explained by the intrinsic properties of the protein or requires other external factors, we expressed and purified recombinant human TagRFP-HURP (**Fig. S3A & B**, see Methods for details). Mixing this protein with dynamic microtubules assembled from GMPCPP stabilized porcine-tubulin seeds showed that at low nanomolar concentrations TagRFP-HURP randomly bound the microtubule lattice but became enriched at the plus-ends of depolymerizing microtubules (**Fig. 3A**), resembling the accumulation of EGFP-HURP at the leading (depolymerization-coupled) kinetochore *in vivo* (**Fig. 1 and 2**). TagRFP-HURP also dramatically decreased the microtubule catastrophe frequency (from 2.93 ± 0.33 h^-1^ for the control to 0.20 ± 0.10 h^-1^ for 50 nM HURP) and increased the rescue frequency (from 282.08 ± 31.15 h^-1^ for the control to 1107.69 ± 553.85 h^-1^ for 50 nM HURP), despite exerting little to no effect on both the polymerization and depolymerisation speeds (**Fig. 3B**). This rescue factor activity of HURP was consistent with its microtubule-stabilizing role in cells (*12*, *14*) and our observed correlation between increasing HURP and the directional switching of the leading kinetochore (**Fig. 2B**).

**Fig. 3.**
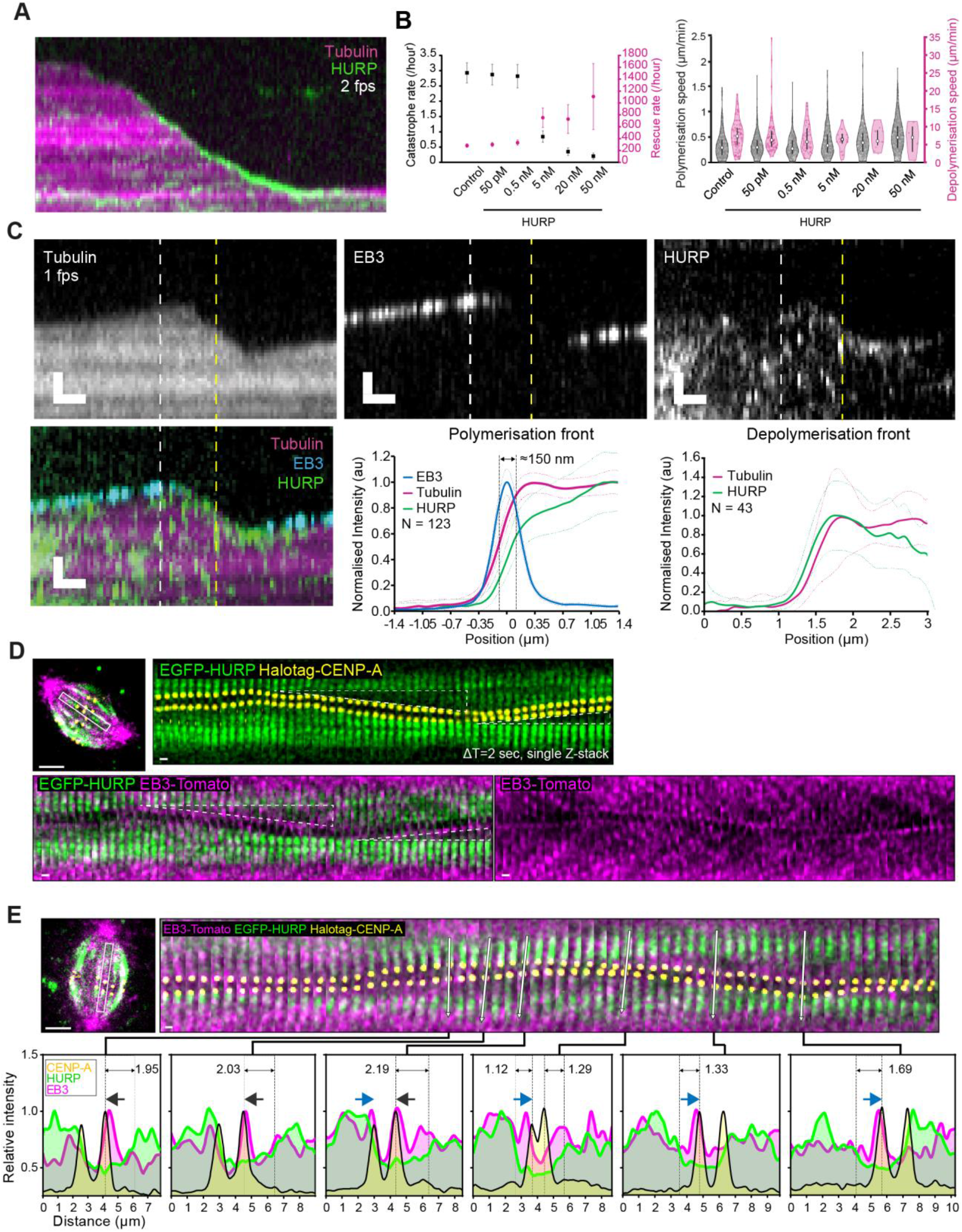
HURP and EB-3 are excluded from a micron-wide zone on growing K-fibres. (**A**) Representative kymograph of 5 nM TagRFP-HURP binding to a dynamic microtubule; scale bar = 1 μm vertically, 10 s horizontally. (**B**) Quantification of the microtubule catastrophe and rescue rate (left), microtubule polymerization rate (middle) and microtubule depolymerization rate (right) at different TagRFP-HURP concentrations (**C**) Representative kymographs of 5 nM TagRFP-HURP and 30 nM EGFP-EB3 binding to a dynamic microtubule (top, scale bar = 1 μm vertically, 10 s horizontally); mean profile of tubulin, EB3 and HURP intensities at the polymerization and depolymerization front (bottom, N = 123 and 43 respectively); white and yellow vertical dashed lines in the kymographs indicate the representative frame used for quantification at the polymerization and depolymerization front, respectively. (**D**) Representative images of HURP, CENP-A and EB3 signals in the hTERT-RPE1 EGFP-HURP/HaloTag-CENP-A cell line overexpressing EB3-Tomato (ΔT = 2 sec). Triangles highlight HURP-gaps. (**E**) Representative intensity line profiles of HURP, CENP-A and EB3 signals based on live cell imaging of hTERT-RPE1 EGFP-HURP/HaloTag-CENP-A overexpressing EB3-Tomato (ΔT = 2 sec). White arrows indicate selected axis of profiling. Dark grey and blue arrows show movement direction of kinetochores. Dashed lines indicate HURP-gaps. (D) and (E); scale bars = 5 μm and 1 μm (kymograph).

We next added EB3-GFP to this minimal system to mark the GTP-cap of polymerising microtubules (*8*, *26*). Kymographs showed that on the microtubule plus-ends TagRFP-HURP mirrors the binding pattern of EB3-GFP, which was only associated with growing plus-ends (**Fig. 3C**). To confirm that recombinant HURP is excluded from the GTP-cap, we averaged the EB3, HURP and tubulin intensity profiles of 123 microtubules on frames where the microtubules were growing. We found that the averaged TagRFP-HURP intensity profile was right-shifted (away from plus-end) by ~150 nm relative to the EB3 peak and the growing microtubule ends. This configuration yielded a gap on single microtubules that contained reduced amounts of EB3 and HURP (**Fig. 3C)** In contrast, such gap could neither be detected on depolymerizing microtubule ends with HURP (**Fig. 3C**), nor was it found at polymerising ends with an alternative end-tracking complex (Ska; **Fig. S3C**). This HURP/EB-depleted zone near the microtubule end was also present in living cells: using hTERT-RPE1 EGFP-HURP/Halo-CENP-A cells expressing EB3-tdTomato we found EB3 closely associated to the trailing kinetochores (bound to growing microtubules). However, when compared to the signal at kinetochores, the EB3 was severely reduced in the HURP-gap on K-fibres (**Fig. 3D and movie 5**). Analysis of the HURP and EB3 intensity profiles revealed an EB3- and HURP-depleted intermediate zone that could reach up to several microns (**Fig. 3E and movie 6**).

Our data indicated that HURP is excluded from GTP cap and prefers the GDP-tubulin lattice both *in vitro* and *in vivo*. This suggested that it may distinguish between the tubulin conformations associated with different nucleotide states. To test this hypothesis we introduced TagRFP-HURP into flow chambers containing surface-bound porcine brain microtubules that were barcoded such that they contained regions with the slowly hydrolysable nucleotide analogues GMPCPP or GTPγS (reported to mimic GDP+Pi or GTP states (*8*, *9*, *27*)) next to regions with GDP (**Fig. 4A**). TagRFP-HURP bound the GDP-tubulin lattice 4-fold higher than the GMPCPP region and ~2-fold higher than the GTPγS-tubulin region (**Fig. 4B**; note that both GMPCPP and GTPγS were pre-treated with taxol, but that its presence did not affect the binding of TagRFP-HURP on those microtubules, **Fig. S4A**). Consistently, TagRFP-HURP molecules had 2-fold higher residency time on the GDP-lattice when compared to the GMPCPP lattice (**Fig. S4B & C**). We conclude that HURP has a preference for GDP microtubule lattice and avoids GTP-like states.

**Fig. 4.**
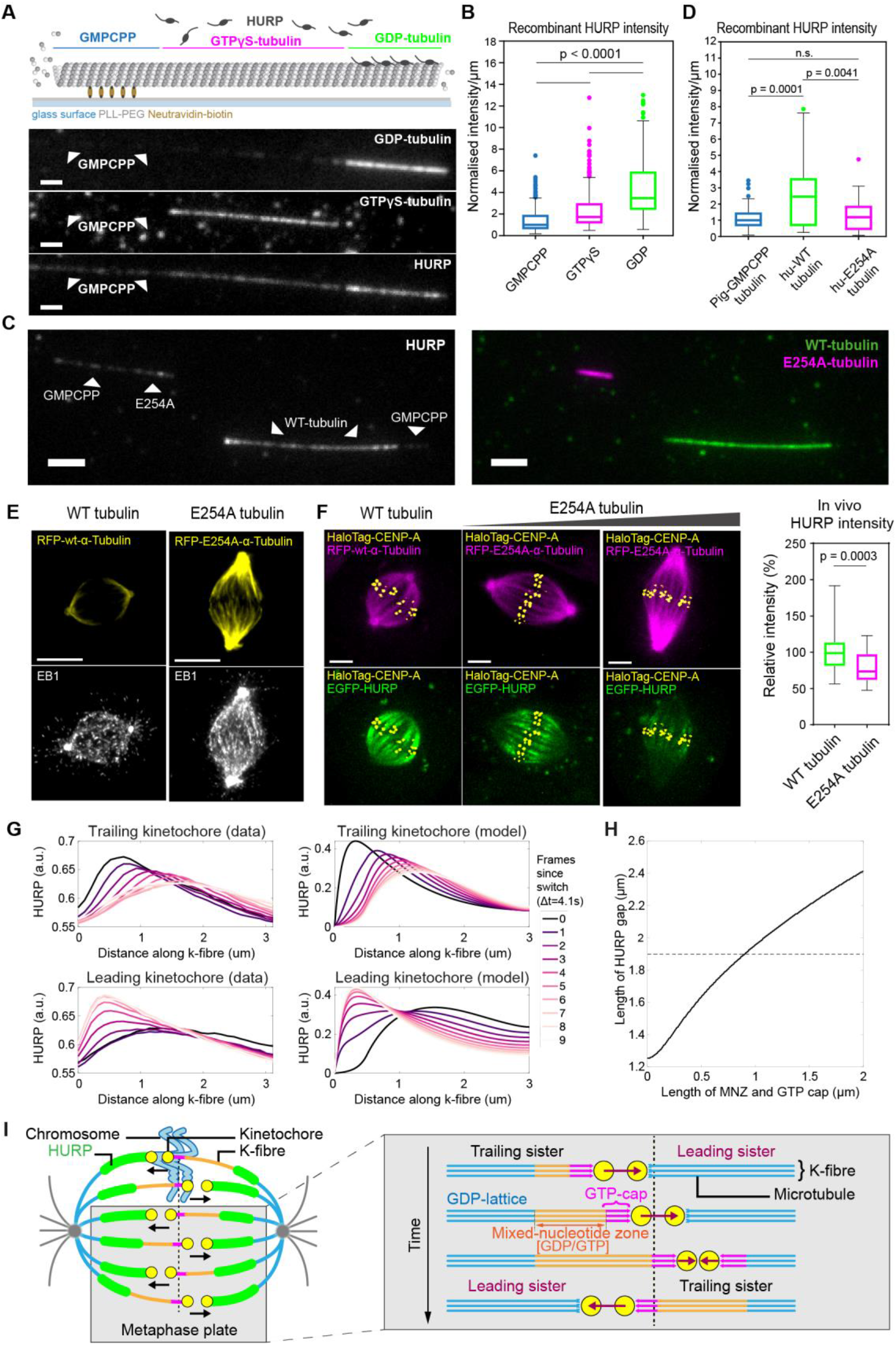
HURP preferentially binds the GDP-lattice of microtubules. (**A**) Schematic of barcoded porcine-tubulin microtubule with the GMPCPP-, GTPγS- and GDP-tubulin sections (top), along with corresponding fluorescence channels of GDP-(Hilyte647), GTPγS-(Hilyte488) tubulin sections (unlabelled GMPCPP-tubulin, see Methods) and TagRFP-HURP (bottom). (**B**) Boxplots of the TagRFP-HURP intensity/μm normalised to its median intensity on GMPCPP microtubules (N = 171 (GMPCPP), 300 (GTPγS) and 238 (GDP); p = Kruskal-Wallis test); scale bars = 1.5 μm. (**C**) Representative image of porcine-tubulin GMPCPP seeds extended with either human WT-(green) or E254A-tubulin (magenta), labelled with Hilyte 488 and Hilyte 647 porcine brain tubulin (1:7 labelled:unlabelled ratio) respectively (right panel) and incubated with TagRFP-HURP (left panel); scale bars = 3 μm. (**D**) Boxplots of HURP intensity/μm normalised to its median intensity on GMPCPP seeds (N = 121 (GMPCPP), N = 43 (WT) and N = 87 (E254A-tubulin); p = Kruskal-Wallis test). (**E**) Immunofluorescence images of metaphase hTERT-RPE1 cells transducted either with RFP-wt-α-Tubulin or RFP-E254A-α-Tubulin and stained for EB1. Scale bars = 5 μm. (**F**) Live cell imaging of hTERT-RPE1 EGFP-HURP/HaloTag-CENP-A cells transducted either with RFP-wt-α-Tubulin or RFP-E254A-α-Tubulin mutant (left), Z-projections of 5 x 0.5 μm, Scale bar = 5 μm; and boxplots of *in vivo* relative HURP intensities (right). N = 2; 38 and 42 cells (p = Mann Whitney test). (**G**) Spatio-temporal dynamics of HURP on K-fibres from *in vivo* data (as in Fig. 2C) and computational model predictions. (**H**) Relationship between length of GTP-cap and mixed-nucleotide zone, and length of HURP gap. (**I**) Proposed model of the mixed-nucleotide zone on K-fibres.

One caveat of this experiment, however, is that these GTP analogues are not necessarily physiological. To substantiate our findings, we also purified wild-type human tubulin and a human tubulin mutant (E254A) incapable of GTP hydrolysis (**Fig. S4D**) (*8*). By extending GMPCPP stabilized porcine-tubulin seeds with either human wild-type- or E254A-tubulin (**Fig. 4C**) we found that TagRFP-HURP avoids the E254A-mutant where the GTP state of tubulin is preserved, but not wild-type human tubulin that hydrolyzes GTP to GDP (**Fig. 4D**). We next considered if this nucleotide dependency also holds in living mitotic cells: we transfected hTERT-RPE1 cells with RFP-tagged wild-type or E254 human tubulin. While both human tubulin variants were well incorporated into mitotic microtubule network, E254A tubulin led to longer mitotic spindles, reduced kinetochore oscillation amplitudes and an increase in the EB1 or EB3 signal (**Fig. 4E and Fig. S4E, movies 7 and 8)**. This indicated an increase in GTP-tubulin within microtubules (*8*). Strikingly, expression of the non-hydrolyzing E254A mutant reduced HURP levels on the mitotic spindle by ~25% (**Fig. 4F and movie 8**). This shows how simply increasing the fraction of GTP-tubulin in the spindle is sufficient to displace HURP molecules. Taken together, our *in vivo* and *in vitro* data reveal how HURP preferentially binds to GDP-tubulin and avoids GTP-tubulin containing microtubule lattice.

Our results reveal the presence of three distinct regions on growing K-fibres: (1) a EB3-positive/HURP-negative GTP-cap abutting the kinetochore, followed by (2) an EB3/HURP-negative zone that can extend for several microns, and (3) a HURP-positive GDP-bound lattice. We postulate that the HURP-gap is caused by a mixed-nucleotide zone that contains insufficient GDP-tubulin or GTP-tubulin to accumulate HURP and EBs respectively. Our *in vitro* data and the fact that E254A tubulin led to the displacement of HURP on the entire spindle, indicate that the HURP-gap is a direct consequence of HURP binding specificity. When simulating HURP dynamics on K-fibres with a minimal mathematical model that incorporates diffusion, exclusion from the GTP-cap and the mixed-nucleotide zone, interaction with the Ran-GTP gradient, and movement of chromosomes, we found that these elements are sufficient to reproduce the observed HURP patterns (**Fig. 4G**). We thus propose that the characteristic HURP-stripes and the associated gaps on growing K-fibres reflect a mixed-nucleotide zone. By plotting the assumed size of the mixed-nucleotide zone in our model versus the size of the HURP-gap, we find that the mixed-nucleotide zone size is smaller than, but proportional to, the HURP-gap size (**Fig. 4H**). Our *in vivo* data show that the transition into and from this mixed-nucleotide zone is sharp for both EB proteins and HURP, suggesting a cooperative binding process for both proteins that requires a specific threshold of GTP- or GDP-tubulin. Consistently, EB proteins bind to microtubules at the interface of 4 tubulin subunits (*28*, *29*) and require GTP-tubulin on adjacent tubulin subunits to bind to microtubules (*26*). We speculate that HURP may recognize the GDP-tubulin conformation at an interface located between adjacent tubulin subunits via a similar mechanism. Future structural analysis will, however, be necessary to dissect the mechanism that confers HURPs preference for GDP-tubulin. Because our *in vitro* data also show that HURP is a potent microtubule rescue factor it is tempting to speculate that HURP stripes establish global boundaries of microtubule depolymerisation lifetime within the spindle. More generally, our work reveals HURP as the first example of a mitotic spindle associated protein that specifically recognizes GDP-tubulin. Since Doublecortin (DCX) and Tau have been shown to also avoid the GTP cap in post-mitotic neuronal cells (*30*, *31*) there is growing support for a new class of growing microtubule tip-avoidance factor.

Our *in vivo* data reveal that the HURP-gap reflects an underlying mixed-nucleotide zone on the growing K-fibre which can reach several microns in cells, much larger than the one detected *in vitro* within a single microtubule (~150 nanometers). This difference could be either due to other microtubule-associated proteins affecting the GTP hydrolysis rate in K-fibres, and/or suggests that parallel bundling of microtubules imparts alterations to the nucleotide and/or structural transitions within the lattices. Low nanomolar concentrations of taxol, which reduces the tubulin-incorporation rate, led to smaller HURP-gaps, implying its size can serve as an indicator for the GTP-tubulin incorporation/hydrolysis rate. More generally, the persistence of the HURP-gap for 45 s and more, indicates that the GTP-tubulin hydrolysis rate in K-fibres is very slow. This suggests that GTP-hydrolysis within the K-fibre associated with the trailing kinetochore is unlikely to dictate sister-kinetochore directional switches. This is consistent with previous observations, showing that such switches are mostly initiated by the leading kinetochore (*32*). In contrast, once a directional switch occurs, HURP-gaps are immediately replenished by HURP, implying a very rapid wave of GTP-hydrolysis along the microtubule lattice once a K-fibre starts shrinking. Overall, the dynamics of HURP indicate that the K-fibre lattice is not a homogeneous structure, but rather that its nucleotide content is highly dynamic in nature. This present work defines the existence of a new mixed-nucleotide zone within the mitotic spindle (**Fig. 4I**) and raises the possibility for equivalent regions in other microtubule-based assemblies.

## Acknowledgments

We thank A. Khodjakov (New York State Department of Health), P. Nowak-Sliwinska (University of Geneva), A. Straube (Warwick University), E. Nigg (University of Basel), E. Dent (University of Madison) D. Gerlich (IMBA, Vienna), M. Strubin (University of Geneva), D. Trono (EPFL, Lausanne) and T. Surrey (CRG, Barcelona) for reagents. We thank the Bioimaging Facility at the Medical Faculty of the University of Geneva and the Computational & Advanced Microscopy Development Unit at Warwick Medical School for microscopy support; N. Liaudet (University of Geneva) for help in data analysis and writing the FRAP analysis code; I. Gasic (University of Geneva) for discussions and advice about fixation methods; A. Diman and M. Strubin laboratory (University of Geneva) for lentiviruses production support; and members of the Meraldi, McAinsh and Gotta (University of Geneva) laboratories for critical discussions.

## Author contributions statement (CRediT)

The original conceptualization of the project was by PM with subsequent ongoing contribution from AM. CC carried out all the cell biology experiments, except Lattice light-sheet microscopy data collection (AM and OS). Image processing, analysis of lattice data and their simulations were carried out by JUH. DD created HURP-EGFP HeLa cell line and made initial HURP dynamics movies. AI carried out all protein purification, *in vitro* reconstitution, and biochemistry experiments. CC and AI wrote an original draft. AM and PM reviewed and edited the manuscript. AM, PM and NB acquired funding and supervised the project.

## Data and materials availability

All data and material can be made available on request.

