## Supplementary material for "Microtubule-based mitotic spindles contain a micron-sized mixed-nucleotide zone"

#### Supplementary figure Legends

#### Figure S1

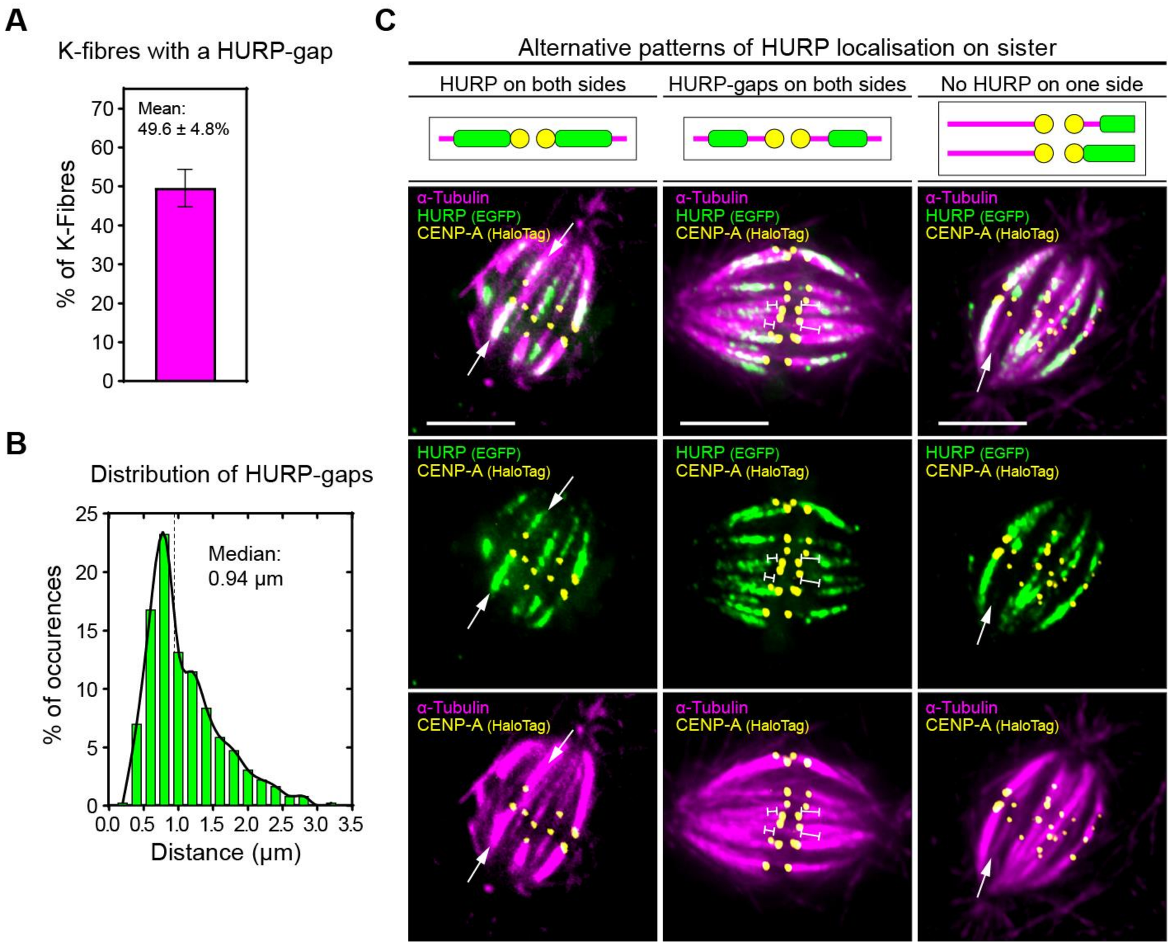


**Fig. Sup 1. Characterization of HURP-gaps and alternative HURP localization patterns on sister K-fibre pairs in fixed cells.** (**A**) Percentage of K-fibres displaying a HURP-gap in hTERT-RPE1 EGFP-HURP/HaloTag-CENP-A cells (error bars = s.d.., N = 4, n = 838 K-fibres in 18 cells). (**B**) HURP-gap length distributions in fixed hTERT-RPE1 EGFP-HURP/HaloTag-CENP-A cells (N = 4, n = 357 gaps in

18 cells). Black line = curve fit. (**C**) Representative images of alternative HURP localization patterns along sister K-fibre pairs observed after fixation of hTERT-RPE1 EGFP-HURP/HaloTag-CENP-A cells. Z-projection of 1.9, 1.6 and 1.9 µm thickness, respectively. Gamma = 0.7 for GFP-channel. Scale bar = 5 µm.

### Figure S2

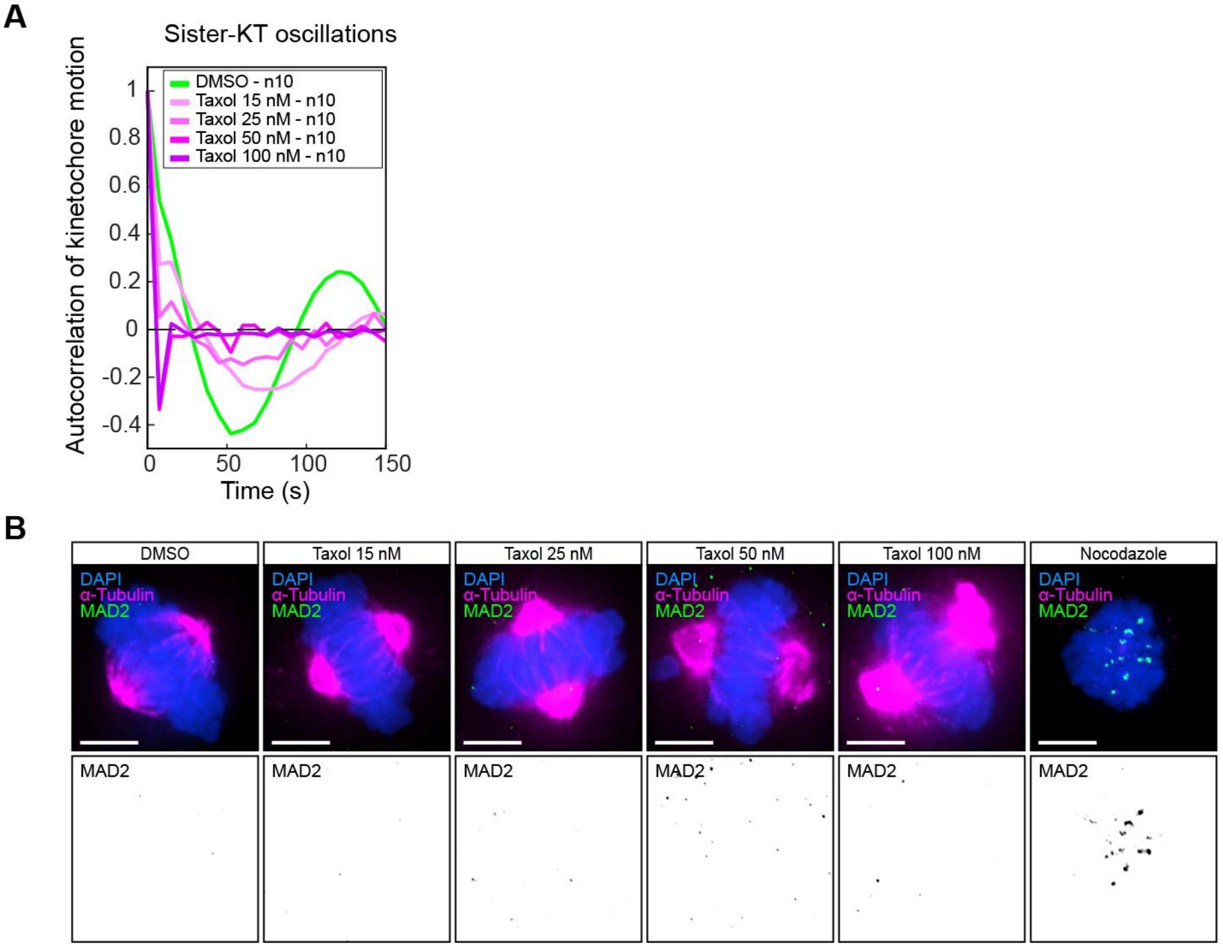


**Fig. Sup 2. Effect of increasing taxol concentrations on kinetochore oscillations and attachment.** (**A**) Autocorrelation curves of sister-kinetochore oscillations in metaphase hTERT-REP1 Centrin1-GFP/GFP-CENPA cells treated with indicated concentrations of taxol for 45 min. (**B**) Immunofluorescence images of metaphase hTERT-RPE1 cells treated with indicated concentrations of taxol for 45 min and stained Mad2. The microtubule depolymerizing drug nocodazole serves as positive control. Scale bar = 5 µm.

### Figure S3

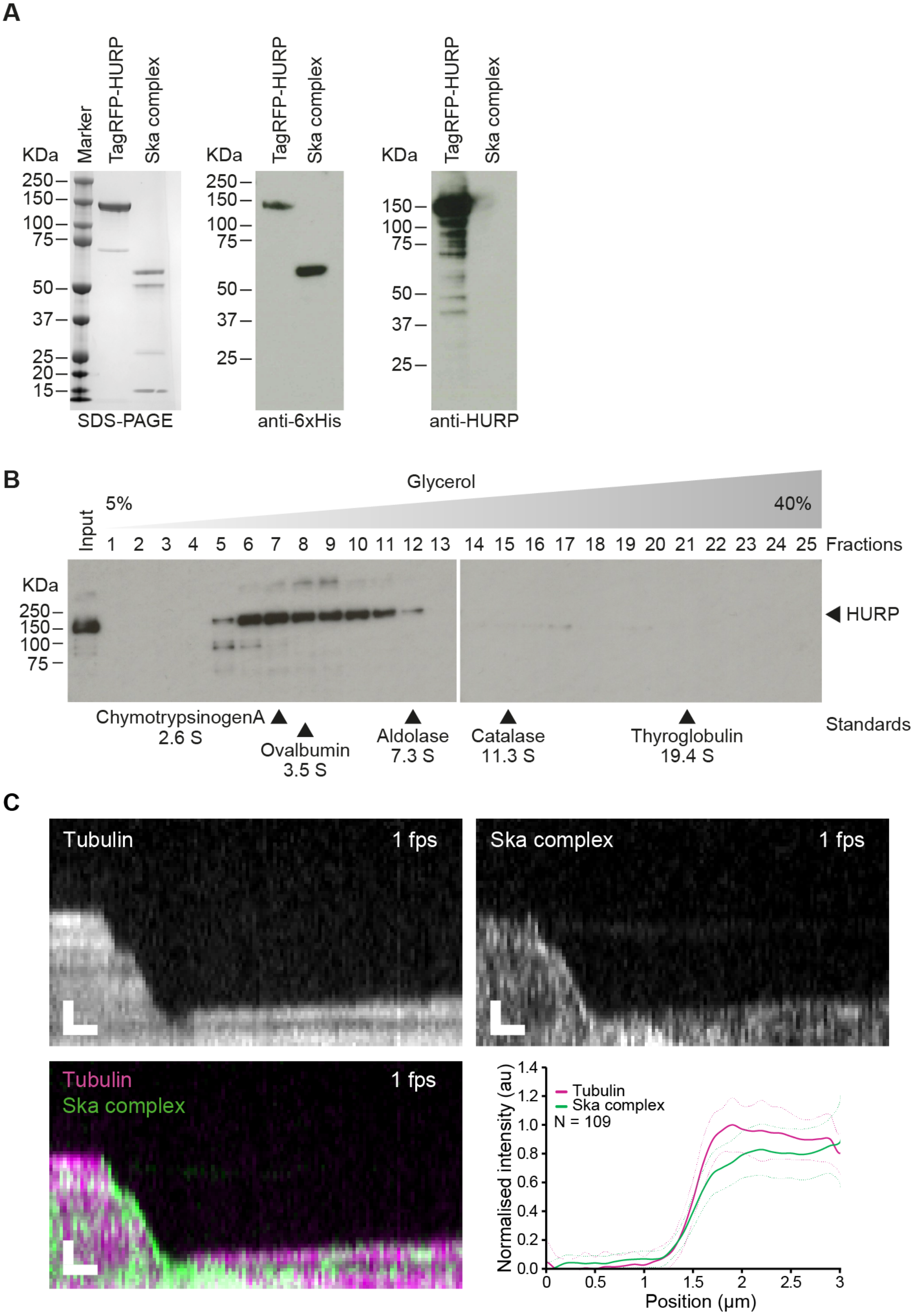

A
KDa
Marker
TagRFP-HURP
Ska complex
250
150
100
75
50
37
25
20
15
SDS-PAGE
anti-6xHis
anti-HURP
B
Glycerol
5%
40%
Input
1 2 3 4 5 6 7 8 9 10 11 12 13 14 15 16 17 18 19 20 21 22 23 24 25 Fractions
HURP
ChymotrypsinogenA 2.6 S
Ovalbumin 3.5 S
Aldolase 7.3 S
Catalase 11.3 S
Thyroglobulin 19.4 S
Standards
C
Tubulin
1 fps
Ska complex
Normalised intensity (au)
Position (µm)
N = 109

**Fig. Sup 3. HURP and Ska complex purification**

(**A**) Purification of TagRFP-HURP and EGFP-Ska complex as shown by SDS-PAGE (left), anti-6xHis (middle, Qiagen penta-His) and anti-HURP (ab70744, Abcam) Western blots. The EGFP-Ska complex is used as a positive (anti-6xHis) and negative (anti-HURP) control. (**B**) Anti-HURP Western blots of 5-40% glycerol gradient fractions, indicating that the purified HURP is monodispersed. (**C**) Representative kymograph of 20 nM EGFP-Ska complex binding to and tracking a depolymerizing microtubule (left) and analysis of its mean intensity profile on the microtubules polymerizing front (N = 109); scale bars are 1 μm vertically, 10 s horizontally.

### Figure S4

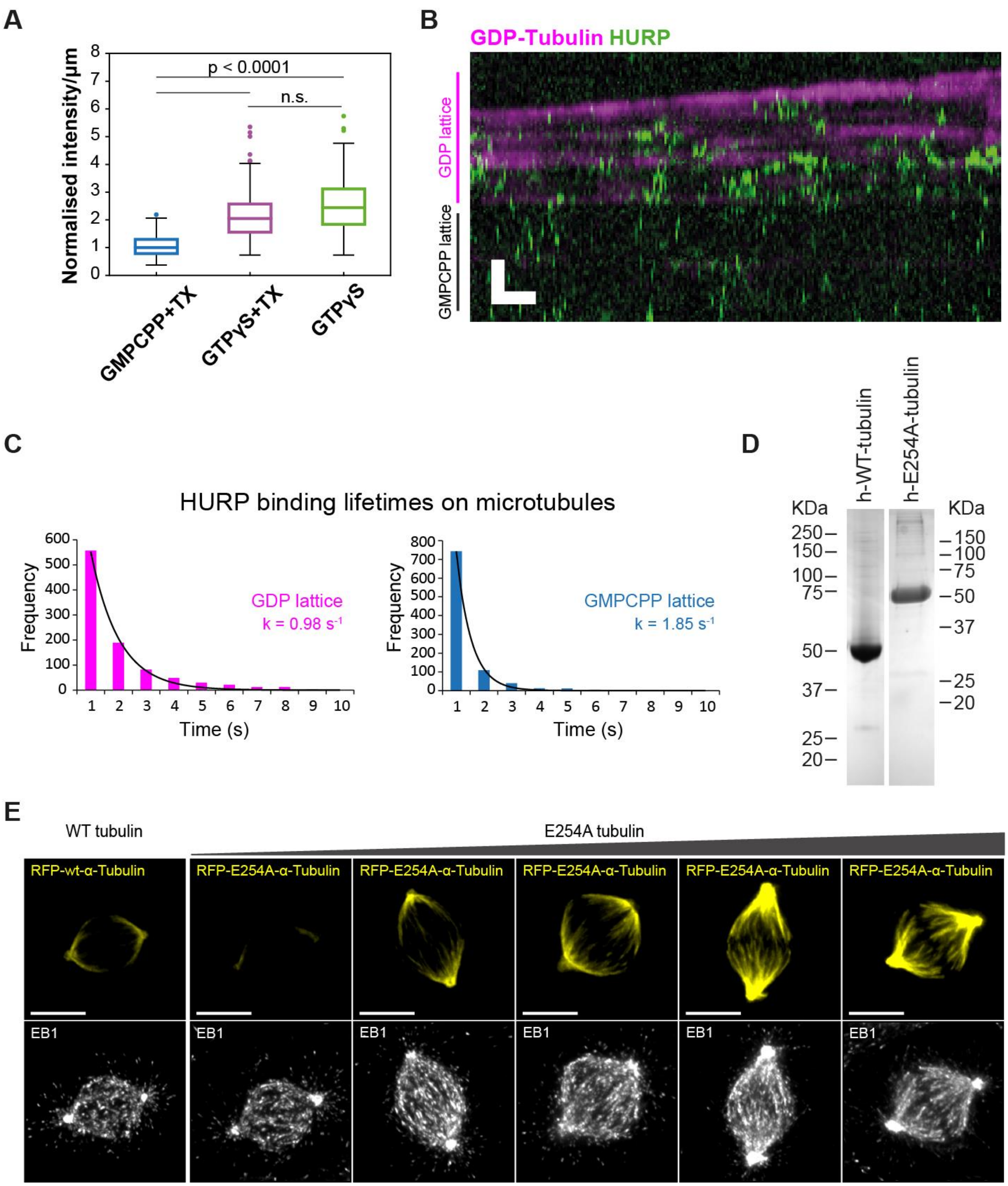


**Fig. Sup 4. HURP preferentially binds GDP-tubulin.** (**A**) Box plots of TagRFP-HURP binding to barcoded microtubules. Note that taxol did not change the amount of TagRFP-HURP bound to GTPγS-tubulin. (**B** and **C**) Representative kymograph of TagRFP-HURP binding on a microtubule (B) and quantification of its lifetimes (C); scale bars = 2 µm vertically; 10 s horizontally. (**D**) SDS-PAGE gels of the purified human WT and E254A tubulin. (**E**) Immunofluorescence images of metaphase hTERT-RPE1 cells transducted either with RFP-wt-α-Tubulin or RFP-E254A-α-Tubulin and stained for EB1, arranged by amount of RFP-E254A-α-Tubulin expression level. Scale bars = 5 µm.

#### Materials and Methods

##### Cell culture, cell lines and treatments.

hTERT-RPE1 (non-transformed immortalized human retina pigment epithelial cells), hTERT-RPE1 Centrin1-GFP/GFP-CENPA (kind gift from A. Khodjakov), HCT-116 (pseudo-diploid colon carcinoma cells; kind gift from P. Nowak-Sliwinska) and hTERT-RPE1 EB3-GFP (kind gift from A. Straube (*26*)) cells were cultured in high glucose DMEM (Thermofisher; 41965) supplemented with 10% FBS and 1% penicillin/streptomycin (Thermofisher; 15140). The medium for hTERT-RPE1 EGFP-HURP/HaloTag-CENP-A, HeLa EGFP-HURP/HaloTag-CENP-A and hTERT-RPE1 EGFP-HURP/HaloTag-CENP-A/EB3-Tomato cells was supplemented with 600 µg/mL of G418 (InvivoGen; ant-gn-5). ECRF24 cells (immortalized human umbilical vein endothelial cells; kind gift from P. Nowak-Sliwinska) were cultured in flasks coated with 0.2% gelatine and grown in 1:1 DMEM-RPMI-1640 (Thermofisher; 41965, 21875) supplemented with 10% FBS and 1% penicillin/streptomycin. To generate the hTERT-RPE1 EGFP-HURP/HaloTag-CENP-A cell line, hTERT-RPE1 EGFP-HURP cells expressing endogenously EGFP-tagged HURP (*14*) were transfected with pHTN Halo-Tag CENP-A. After selection with G418, positive cells for HaloTag-CENP-A were FACS sorted twice. To generate HeLa EGFP-HURP/HaloTag-CENP-A cells, an EGFP tag was introduced by CRISPR/Cas9 into the first exon of the DLGAP5 gene (encoding HURP) in HeLa cells, as previously described (*14*). The selected EGFP positive clone was transfected with pHTN Halo-Tag CENP-A, selected with G418, and sorted for HaloTag-CENP-A expression by FACS twice. To mark kinetochores, cell lines expressing HaloTag-CENP-A were incubated for 30 min with 20 nM of 646 or 549 Janelia Fluor HaloTag ligand (Promega; GA1120, GA1110) 16 hours prior to any experiment. For lattice light sheet imaging, cells were treated with 100 µM TMR (Promega G8252) for 15 min prior to imaging at least 30 min later. The following drugs were used in this study: cells were arrested in metaphase with 10 µM MG132 (Sigma-Aldrich; C2211) for 45 min followed by a treatment with indicated concentrations of taxol (Sigma-Aldrich; T7402) 45 min before acquisition or fixation. Alternatively, 1 µg/mL nocodazole (Sigma-Aldrich; M1404) was used to created unattached kinetochores.

##### Live cell imaging.

Cells were seeded in polymer coverslip 2-, 4- or 8-well µ-Slide ibidi chambers (Ibidi; 80286, 80426, 80826). Cells expressing HaloTag-CENP-A were pre-incubated with HaloTag ligand the day before imaging. Minimum 4 hours prior live cell imaging experiments, the culture medium was replaced by Leibovitz L-15 (Thermofisher; 21083) supplemented with 10% FBS and 1% penicillin/streptomycin. The chambers were acclimatized in a no-$CO_2$ 37°C chamber before imaging. Acquisition was performed using an EC Plan Neofluar 100X (NA 1.3 Oil Ph3) objective on Zeiss Cell Observer.Z1 a spinning disk microscope (Nipkow Disk) equipped with a 37°C chamber, with a HAL100 and HXP120 for transmission and fluorescence widefield visualization and with a CSU X1 automatic Yokogowa spinning disk head. Images of 512 x 512 pixels were acquired with an Evolve EM512 camera (Photometrics) and using Visiview 4.00.10 software. For *in vivo* visualization and quantification of EGFP-HURP and HURP-gaps on K-fibres, metaphase hTERT-RPE1 EGFP-HURP/HaloTag-CENP-A or HeLa EGFP-HURP/HaloTag-CENP-A cells were imaged every 5 s for 2 or 3 min with 5 Z-stacks of 0.5 µm spacing. For co-visualization of EGFP-HURP and EB3-Tomato on K-fibres or visualization of EB3-GFP in presence of wt- or E254A-α-Tubulin, metaphase or interphase cells were imaged on a single focal plane or alternatively with 3 Z-stacks of 0.5 µm spacing, every 2 s for 2 min. Lattice light sheet imaging was performed with a lattice light sheet microscope from Intelligent Imaging Innovations (3i; *https://www.intelligent-imaging.com*). Cells were seeded on 5 mm radius glass coverslips one day before transfer of coverslip to the LLSM bath filled with $CO_2$-independent L15 medium, where live imaging takes place. 3D time-lapse images (movies) of hTERT-RPE1 EGFP-HURP/HaloTag-CENP-A cell line were acquired at 488 nm and 560 nm channels using 1% and 4% laser power respectively, 35 ms exposure time/z-plane, 90 z-planes, 308 nm z-step, which results in 4.14 s/z-stack time (frame). Acquired movies were de-skewed and cropped in XYZ and time, using Slidebook software. Cropped movies were then saved as OME-TIFF files in ImageJ.

**Immunofluorescence.**

For visualization of EGFP-HURP or endogenous HURP and quantification of gaps on K-fibres, cells were grown on acid-etched glass coversplis, permeabilized with 0.5 % Triton X-100 in 37°C pre-warmed Microtubule Stabilizing Buffer (MTSB buffer – 80 mM KOH-PIPES (pH = 6.8), 1 mM $MgCl_2$, 5 mM EGTA) for 30 s and fixed with 0.5% Glutaraldehyde (Sigma-Aldrich; G6403) in MTSB buffer for 10 min at RT. After fixation, glutaraldehyde was quenched by 0.1% $NaBH_4$ in PBS for 7 min at RT, followed by 2 x 3 min PBS washes. Cells were blocked for 30 min in PBS + 3% BSA and immunolabelled. For cells expressing EGFP-HURP, anti-GFP and anti-α-tubulin antibodies were added for 1 h at RT followed by 3 x 3 min PBS washes before a 1 h incubation with appropriate Alexa Fluor–conjugated secondary antibodies (1:400; Invitrogen). Immunolabelled cells were washed 3x with PBS before mounting the coverslips with VECTASHIELD with/without DAPI (Vector Laboratories). For detection of endogenous HURP in hTERT-RPE1, HCT-116 and ECRF24 cell lines, the anti-HURP, anti-Hec1 and anti-α-tubulin primary antibodies were added for 2 h.

For detection of Mad2, cells were fixed 10 min with 4% formaldehyde (Sigma-Aldrich; 47608) in 20 mM KOH-PIPES (pH = 6.8), 10 mM EGTA, 1 mM $MgCl_2$, 0.2% Triton X-100, rinsed 2 x 3 min with PBS, blocked for 30 min in PBS + 3% BSA and stained. For visualization of EB1 cells were fixed in -20°C methanol (Sigma-Aldrich; 32213) for 6 min, rinsed 3 min twice with PBS, blocked for 30 min in PBS + 3% BSA and stained. All primary and secondary antibodies were diluted in PBS + 3% BSA.

The following primary antibodies were used: rabbit anti-GFP (1:500; (*33*)), chicken anti-GFP (1:1000; Thermofisher; A10262), rabbit anti-HURP (1:500; kind gift from E. Nigg; (*12*)), recombinant human anti-α-tubulin (1:300; (*34*)), rabbit anti-α-tubulin (1:1000; abcam; ab18251), mouse anti-Hec1 (1:700; abcam; ab3613), rabbit anti-Mad2 (1:1000; Bethyl; A300-301A) and mouse anti-EB1 (1:200; BD biosciences; 610535).

Immunofluorescence images were acquired on an Olympus DeltaVision wide-field microscope (GE Healthcare) equipped with a DAPI/FITC/TRITC/Cy5 filter set (Chroma Technology Corp.) and a Coolsnap HQ2 CCD camera (Roper Scientific) running Softworx (GE Healthcare). 3D images were deconvolved using Softworx (GE Healthcare). To visualize EGFP-HURP or endogenous HURP and quantify HURP-gaps, mitotic spindles were imaged using a 100X 1.4 NA objective in 0.1 µm Z-stacks; to visualize Mad2, mitotic spindles were imaged using a 100X 1.4 NA objective in 0.2 µm Z-stacks; to visualize EB1, mitotic spindles and interphase cells were imaged with a 100X 1.4 NA or a 60X NA 1.4 objective in 0.2 µm Z-stacks. Endogenous HURP and HURP-gaps in hTERT-RPE1 cells were visualized with an EC Plan Neofluar 100X (NA 1.3 Oil Ph3) objective in 0.1 µm Z-stacks on a spinning disk microscope (Nipkow Disk) Zeiss Cell Observer.Z1 equipped with a HXP120 fluorescence widefield visualization lamp and with a CSU X1 automatic Yokogowa spinning disk head. Images of 512 x 512 pixels were acquired with an Evolve EM512 camera (Photometrics) and the Visiview 4.00.10 software.

**Fluorescence recovery after photobleaching.**

Cells were seeded in glass bottom 4-well µ-Slide 4 Ibidi chamber (Ibidi; 80427). One day before acquisition, hTERT-RPE1 EGFP-HURP/HaloTag-CENP-A cells were incubated for 30 min with 20 nM (1:10 000) of Janelia Fluor 646 HaloTag Ligand to mark kinetochores. Cells were incubated in imaging medium 4 hours prior FRAP experiment. Metaphase cells were selected based on brightfield contrast and a single focal planes of 140 nm pixel size were acquired using a 60X (NA 1.4) CFI Plan Apochromat oil objective on a Nikon A1r point scanning confocal microscope and NIS elements software. Single K-fibres were visualized based on EGFP-HURP signal and targeted with a circular ROI of 10 pixels (1.3 µm) in diameter before photobleaching with two 500 ms pulses of 80% 405 nm laser power. After bleaching, the cells were acquired on single focal plane every 2 s for 2 min.

FRAP experiments were analyzed using a custom-made application written in Matlab 2021a (MathWorks, Natick MA USA), *code is available on GitHub https://github.com/Bioimaging/FRAP*. Briefly, if present, time lapse images drift was corrected using rigid registration. Three classes of region

of interest were manually positioned to report the bleached area, the whole mitotic spindle and the background area fluorescent intensities.
The FRAP signal was computed using a double normalization procedure as followed:

$$FRAP(t) = \frac{Bleach(t) - Back(t)}{Bleach_{pre} - Back_{pre}} \cdot \frac{Ref_{pre} - Back_{pre}}{Ref(t) - Back(t)}$$

Where Bleach(t) is the spatial average intensity in the bleached region, $Bleach_{pre}$ is the average intensity before the bleaching pulse; Ref(t) is the spatial average intensity in the whole mitotic spindle, $Ref_{pre}$ is the average intensity before the bleaching pulse; finally, Back(t) is the spatial average intensity in the background region, $Back_{pre}$ is the average intensity before the bleaching pulse. Such a normalization allows among other things to account for the photobleaching induced by the imaging laser.

**Image processing and analysis.**
To quantify the maximum distance and duration of HURP-gaps in live cell imaging data, the acquired movies were visualized and analyzed using Imaris 7.7 software (BitPlane). A maximum intensity projection (Z-projection) was first performed to obtain 2D movies and if required, over-time translational and rotational drift correction was applied using surfaces tool based on whole EGFP-HURP signal. Kinetochores were automatically or manually detected using spots tool on CENP-A signal. To quantify the maximum duration of HURP-gaps on selected K-fibres, the numbers of time frame (every 5 s) from starting growing K-Fibre until the next directional switch of kinetochores were counted.
The maximum distance of HURP-gaps was measured using the Measurement points tool and by taking the distance between the centre of CENP-A signal (kinetochore) to the edge of the EGFP-HURP signal on growing K-Fibres on the time frame just before the next directional switch.
To visualize HURP localization on sister K-fibre pairs and quantify HURP-gap distances in fixed cells, deconvolved images were visualized in 3D using Imaris 7.7. Only cells displaying parallel mitotic spindles to x-axis were selected and sister K-fibre pairs were identified based on α-tubulin and CENP-A signals. Gamma for GFP/HURP-channel was adjusted to increase visibility of HURP-gaps, as indicated in figure legend. The Ortho Slicer tool was used on XY plane to browse stack by stack in 3D the whole mitotic spindle. To measure HURP-gap distances, the distance between the centre of the CENP-A signal to the EGFP-HURP edge on the same K-fibres were measured using the Measurement points tool.
EGFP-HURP intensities on mitotic spindles of immunolabelled taxol-treated hTERT-RPE1 EGFP-HURP/HaloTag-CENP-A cells were quantified using ImageJ/Fiji software. Due to high cytosolic EGFP-HURP background in presence of taxol, EGFP-HURP intensity was measured in 2D on three different slices (Z-stack) localized on upper, middle, and lower parts of the mitotic spindle. To measure EGFP-HURP intensity, the integrated density (IntDen) of EGFP-HURP signal was measured on half mitotic spindle of each selected slices, inside an ROI of 25 x 120 pixels (1.620 µm x 7.776 µm) enclosing K-fibres close to kinetochores (CENP-A signal). EGFP-HURP background intensity was then determined by moving ROI in a cytosolic region close to the spindle pole. After background subtraction, the three EGFP-HURP intensities were averaged.
EGFP-HURP intensities in live cell imaging movies of hTERT-RPE1 EGFP-HURP/HaloTag-CENP-A cells transfected with WT- or E254A-α-Tubulin were quantified using ImageJ/Fiji software. Z-projection of movies (5 x 0.5 µm stacks) was first applied using the Sum slices tool and the integrated density (IntDen) of EGFP-HURP signal was measured on the 3$^{rd}$ frame of imaging (after 10 s). Measurement was performed with an ROI of 50 x 100 pixels (8 µm x 16 µm) enclosing K-fibres (RFP signal) of the kinetochore region (CENP-A signal). The EGFP-HURP background intensity was measured with an ROI of 30 x 30 pixels (4.8 µm x 4.8 µm) inside the cytosol, before subtracting it from the EGFP-HURP intensity on the mitotic spindle.

**Kinetochore tracking assay** hTERT-RPE1 Centrin1-GFP/GFP-CENPA cells were seeded in a 4-well µ-Slide ibidi chambers (Ibidi; 80426), treated with 10 µM of MG132 (Sigma-Aldrich; C2211) for 45 min followed by a treatment with indicated doses of taxol (Sigma-Aldrich; T7402) for additional 45 min before acquisition. Metaphase arrested cells were imaged in the GFP channel with 2 x 2 binning, every 7.5 s for 5 min and 15 µm Z-stack were acquired every 0.5 µm spacing. All movies were acquired using a 100x NA 1.4 objective on an Olympus DeltaVision wide-field microscope (GE Healthcare) equipped with an eGFP/RFP filter set (Chroma), and a Coolsnap HQ2 CCD camera (Roper Scientific) running Softworx software (GE Healthcare). 3D movies were deconvolved and cropped using Softworx software (GE Healthcare). Deconvolved movies (4D images XYZT) were analyzed using automated kinetochore tracking software (KiT) written in MATLAB 2013b (MathWorks; (*35*)). The latest version of the MATLAB code is available on *https://github.com/cmcb-warwick* and is described in Olziersky et al, (*36*). Briefly, the frame-to-frame displacement of sister-kinetochores and their relative distance from the center of the metaphase plate are analyzed and an autocorrelation function is used to quantify the regularity of the sister-kinetochore oscillations along the spindle axis. For lattice light sheet imaging, kinetochore tracking and sister pairing was performed using the CENP-A channel with a version of the kinetochore tracking software (KiT) adapted for lattice light sheet data (*17*). Line profiles were measured in the direction of the inter-kinetochore vector (smoothed in time with a 5 point stencil) to quantify the spatial distribution of HURP on K-fibres. Line profiles extend 3 µm along the K-fibre and linear interpolation between pixels is applied where necessary. K-fibre polymerisation states were annotated automatically via Bayesian inference using a model of kinetochore dynamics in metaphase as in previous work (*16*). Directional switches from a leading to trailing state were identified in each MCMC draw, and averages taken over such coherent switch events and MCMC draws to provide the average spatial profile of HURP along K-fibres.

**Plasmids and transfection.**

To generate the pHTN Halo-Tag CENP-A plasmid, a synthetic cDNA encoding human CENP-A (Histone H3-like centromeric protein A) was cloned into pHTN HaloTag CMV-neo vector (Promega) using EcoRI and XbaI restrictions sites. The EB3-tdTomato plasmid (kind gift from E. Dent; Addgene 50708) was used to follow GTP caps of microtubules. For transfections, the culture medium was exchanged with MEM (Thermofisher; 41090) supplemented with 10% FBS and 1% penicillin/streptomycin (Thermofisher; 15140) prior transfection. Cells were transfected with X-tremeGENE 9 DNA Transfection Reagent (Roche) with a ratio of 3:1 µl/µg DNA according to the manufacturer's instructions. The TagRFP-HURP construct used for all *in vitro* experiments was ordered from GeneArt using the HURP gene sequence for isoform 1 (Uniprot Q15398), optimised for Sf9 insect cell expression (Spodoptera Frugiperda). The construct starts at the N-terminus with a 6x Histidines tag, followed by a linker containing a Prescission protease recognition sequence (SGVLFQGP) and a NotI and NdeI restriction site, before the start of the TagRFP sequence. This is further separated from the HURP sequence by a linker comprising of five Glycines surrounded by an XhoI and NdeI restriction sites.

**Lentivirus production and transduction of RFP-wt- or RFP-E254A-α-tubulin.**

The α-tubulin E254A mutant was generated by site-directed mutagenesis (Stratagene) using the pmRFP alpha tubulin IRES puro 2b plasmid as template (kind gift from D. Gerlich; Addgene 21041). For subcloning, the pCMV WT- and E254 RFP alpha tubulin sequence were introduced into the pCDH-CMV hGLuc (kind gift from M. Strubin) receiver plasmids using the SpeI and NotI restrictions sites.

For production of VSV-G pseudotyped recombinant lentiviruses, $4.5 \times 10^6$ HEK 293T/17 cells (ATCC; CRL-11268) were seeded in a 10-cm dish and transiently transfected for 16 h using the calcium phosphate method with 10 µg of packaging plasmid psPAX2 (Kind gift from D. Trono; Addgene 12260), 5 µg of envelope plasmid pMD2G (kind gift from D. Trono; Addgene 12259) and with 15 µg of pCDH-CMV-RFP-alpha-Tubulin (wt) or pCDH-CMV-RFP-alpha-Tubulin E254A mutant vector. The medium was replaced 8 h after transfection. Supernatants containing recombinant viruses were collected 48 h post-transfection and filtered through PVDF 0.45 µm filters (Merck-Millipore). Recombinant viruses were stored at -80°C until use.

For lentiviral transduction, $2 \times 10^4$ cells/well were seeded in a 4-well µ-Slide ibidi chambers (Ibidi; 80426). The day after, cells were incubated with different volume of virus containing supernatant. 24 h later, cells were washed several times with PBS and incubated with Janelia Fluor 646 HaloTag ligand before being placed in fresh medium for additional 24 h before imaging.

**Protein purification**

6xHis-TagRFP-HURP was expressed in Sf9 insect cells. Pellets from 2 L cultures were resuspended in Lysis buffer (50 mM Hepes pH 7.5, 200 mM NaCl supplemented with 1% Igepal, 5% glycerol, 10 mM $MgCl_2$ and DnaseI) and the lysate homogenised with 10 strokes, supplemented with 5 mM DTT, 1% PMSF and 1% SERVA antiprotease and incubated on ice for 20 min. The lysate was sonicated and centrifuged at > 40000 g for 20 min at 4°C, and the supernatants loaded onto a SP FF 5 ml column (Cytiva) equilibrated in Lysis buffer. The column was washed with Lysis buffer, followed by Buffer A (50 mM Hepes pH 7.5, 240 mM NaCl, 5 mM DTT) and eluted via a 20 CV gradient from 5 to 100% of Buffer B (50 mM Hepes pH 7.5, 1 M NaCl, 5 mM DTT). The eluted fractions were pooled and concentrated into a 30 kDa MWCO spin column (Amicon) at 3240 g for 40 min at 4°C, diluted 7-fold using Buffer A and loaded onto two serially connected 1 ml Q sepharose FF columns equilibrated in Buffer C. These were washed with 8 CV of Buffer C, then both flowthrough and washes were collected, pooled and concentrated as above to exchange in Buffer C (50 mM Hepes pH 7.5, 200 mM NaCl, 10 mM Imidazole, 1 mM DTT and 0.05% Tween20). The sample was loaded onto a 5 ml HisTrap FF (Cytiva) equilibrated in Buffer C. The column was washed with 8 CV of Buffer C and eluted with a 10 CV gradient to 100% of Buffer D (50 mM Hepes pH 7.5, 200 mM NaCl, 500 mM Imidazole, 1 mM DTT and 0.05% Tween20). The isolated fractions were pooled and concentrated as above and subsequently injected into a Superose 6 Increase gel filtration column (Cytiva) equilibrated and ran in Buffer E (50 mM Hepes pH 7.5, 300 mM NaCl, 1 mM DTT). Finally, fractions were pooled, aliquoted and flash frozen and stored in liquid nitrogen.

The human tubulin wt- and E254A constructs were a kind gift by T. Surrey and were expressed in Sf9 insect cells and purified as described in Roostalu et al, 2019 (*8*), while the EB3-GFP purified protein was kindly gifted by A. Straube. Finally, the EGFP-Ska complex was expressed and purified as in Maciejowski et al, 2017 (*37*).

**Imaging HURP *in vitro***

The *in vitro* microtubule assay was performed as described in Maciejowski et al, 2017 (*37*). In brief, microtubules seeds stabilised with 1 mM GMPCPP and 2 µM taxol were adhered to the coverslip surface of a flow chamber via biotin-streptavidin link. After incubating for 5 min, the chamber was perfused with the assay mix, comprising of 9 µM free tubulin (1:25 unlabelled:labelled ratio using either Hilyte647 or Hilyte488, Cytoskeleton) in BRB80 buffer (80 mM K-PIPES pH 6.8, 1 mM MgCl2, 1 mM EGTA) supplemented with 70 mM KCl and Oxygen scavengers (0.4 mg/ml glucose oxidase, 0.2 mg/ml catalase, 50 mM glucose) and the protein of interest. This allows us to study the binding and unbinding of the protein of interest while the microtubules are undergoing dynamic instability.

Barcoded microtubules were created by adding 16 µM of pig brain tubulin (1:15 labelled:unlabelled) to stabilised GMPCPP seeds (final ratio of 1:1 v:v), in the presence of 1 mM GTPγS, and incubating the solution at 37°C for 1 hour. The mix was then diluted 5 folds in BRB80 supplemented with 2 µM taxol (BRB80+Tx), spun in an airfuge at 20 psi for 10 min at room temperature and the pellet finally resuspended in BRB80+Tx. For the human tubulin experiments, 16 µM of wt-tubulin or 6 µM of E254A-tubulin were added to GMPCPP seeds (final ratio of 5:1 v:v), in the presence of 1 mM GTP.

Chambers were imaged in TIRF while maintaining a constant temperature of either 30°C, when looking at the effect of TagRFP-HURP on microtubules dynamics (Fig. 3B) and when looking at the binding properties of TagRFP-HURP on barcoded microtubules (Fig. 4), or 35°C, when investigating the EGFP-EB3/TagRFP-HURP binding in *vitro* (Fig. 3A and C).

**Minimal model of HURP dynamics on K-fibres**

In the minimal model of HURP dynamics, we account for diffusion of HURP, preferential binding of HURP to the GDP lattice and exclusion of HURP from the GTP cap, effects of the Ran GTP gradient on HURP binding dynamics, and movement of the chromosomes. We take a co-ordinate system along the K-fibre adjacent to the kinetochore in the direction of the spindle pole. The model takes the form of a continuum partial differential equation with advection, diffusion and reaction terms as follows:

$$\frac{\partial H}{\partial t} + v\frac{\partial H}{\partial x} = D\frac{\partial^2 H}{\partial x^2} + \lambda(x)g(x) - \mu H,$$

where $H$ is the concentration of HURP, $D$ is the diffusion coefficient of HURP, $\lambda(x)$ is the binding rate, $g(x)$ is the RanGTP gradient, $\mu$ is the unbinding rate, and $v$ is the speed of chromosome directional movements. We assume no flux boundary conditions at the spindle pole, and assume no HURP at the kinetochore giving a Dirichlet condition at 0. We assume that HURP is excluded from the GTP-cap and the mixed nucleotide-zone, meaning that $\lambda(x)$ takes the form:

$$\lambda(x) = \begin{cases} \lambda_{MNZ} for x \in [0, l), \\ \lambda_{GDP} for x \in [l, L], \end{cases}$$

where $l$ is the length of the GTP-cap and mixed nucleotide-zone. For leading kinetochores, we can assume $l = 0$ and only the dynamics on the GDP lattice are relevant. We assume that the RanGTP gradient is stationary and can be described by $g(x) = exp(-x/s)$ where $s$ is a spatial scale for the RanGTP gradient. Thus the only difference in the model between leading and trailing kinetochores is the exclusion of HURP in the mixed nucleotide zone for trailing kinetochores, and movement of chromosomes in opposite directions relative to the coordinate system. To provide initial conditions, the system is simulated up to equilibrium for the trailing kinetochore, and this equilibrium distribution is used as the initial condition for the leading kinetochore, which is in turn used as the initial condition for simulating the trailing kinetochore. The model is simulated numerically with parameter values given in Table S1 using the MATLAB partial differential equation solver, pdepe.

**Statistical analysis.**
Statistical analysis for figure 2H and 4D were performed using GraphPad Prism 8 (GraphPad), the statistical tests employed are described in the figure legends. Analysis of the intensities of HURP and the Ska complex in all *in vitro* experiments was performed using Excel, while all p values were obtained using Matlab.

Table S1

| Parameter | Value | Units | Description | Reference |
|---|---|---|---|---|
| L | 3 | μm | Size of domain | - |
| T | 37 | s | Final simulation time | - |
| l | 0.5 | μm | Size of GTP-cap and mixed-nucleotide zone | - |
| s | 1 | μm | Spatial scale of RanGTP gradient | 38 |
| v | ± 0.03 | μm/s | Velocity of trailing/leading kinetochore | 16 |
| D | 0.005 | $\mu m^2/s$ | Diffusion constant along K-fibre | - |
| μ | 0.05 | $s^{-1}$ | HURP unbinding rate | - |
| $\lambda_{GDP}$ | 0.05 | $s^{-1}$ | HURP binding rate on GDP lattice | - |
| $\lambda_{MNZ}$ | 0 | $s^{-1}$ | HURP binding rate on GTP-cap and mixed-nucleotide zone | - |
